# Assessing the effectiveness of honey bee pollinators for cultivated blueberries in South Africa

**DOI:** 10.1101/2021.09.03.458826

**Authors:** Keanu Martin, Bruce Anderson, Corneile Minnaar, Marinus L de Jager

**Affiliations:** Department of Botany and Zoology, Stellenbosch University, Matieland 7602, Cape Town, Western Cape, South Africa

**Keywords:** Honey bee, blueberries, pollination, agriculture, pollination ecosystem service, yield

## Abstract

Globally, agricultural crops are often dependent on insect pollination. Blueberries are an example of such a crop and owing to their proposed health benefits they are grown around the world, including locations where their native bumble bee pollinators do not occur. In the absence of bumble bees, blueberry pollination in South Africa and many other parts of the non-native, commercial range is performed primarily by honey bees. Despite this, the effectiveness of honey bee pollination on blueberries remains understudied. This study determined the effect of honey bee pollination on components of fruit yield (fruit set and mass) of five blueberry varieties that are extensively planted in South Africa. For each variety, two metrics were calculated: 1) the benefit of bees — a comparison of fruit yields after exposure to honey bees and fruit yields after honey bee exclusion, 2) the pollination deficit — the difference in yield between hand pollination (maximum yield potential) and yields after exposure to honey bee pollinators. Honey bee pollination consistently resulted in improved yields, although the magnitude of this improvement (i.e., the benefit of bees) was dependent on the variety considered. Similarly, the pollination deficit also varied considerably across varieties and while some varieties appeared to perform close to maximum potential (small pollination deficit), others yielded well below their maximum potential under honey bee pollination. This study demonstrates that honey bees are functional pollinators of blueberries in areas where native blueberry pollinators are absent. However, in such areas, it is important that special focus be given to selecting blueberry varieties that perform well with honey bees as their sole pollinator. Further research is necessary to determine how the pollination deficit of blueberry varieties can be decreased as well as how to increase the effectiveness of honey bee pollination.

## 1. Introduction

Insect pollination is vital for numerous crops, not only to set fruit but also to increase fruit yield (Klein et al., 2007). Consequently, this ecosystem service has been valued at €153 billion per year (Gallai et al., 2009). Although honey bees are globally the most widely used commercially managed pollinator (Garibaldi et al., 2013; Osterman et al., 2021), they are not always the most efficient pollinators and can be outperformed by wild insects (Garibaldi et al., 2013; Osterman et al., 2021). In a comprehensive study using 41 crops, Garibaldi et al. (2013) found that wild insects were often more effective crop pollinators than honey bees and that on average, fruit set doubled after increasing wild insect visitation compared to equivalent increases in honey bee visitation.

One crop where honey bees may be less effective than other pollinators is blueberries. Wild and cultivated blueberries are extremely reliant on insect pollination (MacKenzie, 1997; Fulton et al., 2015; Reilly et al., 2020; Martin et al., 2021;) and are typically pollinated by large bees. These include blueberry bees (Apidae) (Cane and Payne, 1993; Campbell et al., 2018), bumble bees (Apidae) (Mackenzie and Eickwort, 1996; Stubbs and Drummond, 2001; Javorek et al., 2002; Tuell et al., 2009; Campbell et al., 2017), and mining bees (Andrenidae) (Mackenzie and Eickwort, 1996; Javorek et al., 2002). These bees are thought to be effective pollinators of blueberry flowers due to their ability to buzz pollinate which causes the poricidal anthers to release their pollen (Cane and Payne, 1993). In comparison, honey bees cannot buzz pollinate and are less effective at releasing pollen from these specialised anthers. Consequently, bumble bees deposit about two-thirds more pollen per visit than honey bees (Javorek et al., 2002). However, recently Hoffmann et al. (2018) revealed that honey bees that foraged on blueberries had blueberry pollen on all body parts with the majority of blueberry pollen being found on their legs and tarsi. These pollen-laden legs and tarsi regularly came into contact with blueberry stigmas during landing, grooming, and foraging and have the ability to effectively pollinate blueberry flowers. Moreover, Martin et al. (2021) showed that honey bees effectively pollinate Ventura, a blueberry variety commonly grown in South Africa, by significantly increasing both blueberry mass and diameter, as well as decreasing ripening time.

Despite these promising results, the perceived inefficiency of honey bees as pollinators has resulted in pressure to import native blueberry pollinators, such as bumble bees. This is extremely problematic as introducing bumble bees into novel areas has the potential to disrupt local ecosystems. For example, introduced bumble bees can outcompete native pollinators, damage flowers, and rob nectar (Morales et al., 2013; Schmid-Hempel et al., 2014; Aizen et al., 2018). Consequently, it is important to quantify the benefits of honey bees for different commercially planted blueberry varieties so that objective decisions can be made about the perceived needs of importing novel pollinators. This may also enable us to farm more intelligently with varieties that do better under honey bee pollination.

We aim to determine the effects of honey bee pollination on the fruit characteristics of five blueberry varieties that are extensively planted in South Africa. Particularly, we compare the benefits in terms of fruit yield (fruit mass, fruit diameter, percentage fruit set, and adjusted fruit mass) after honey bee pollination with fruit yield when pollinators are absent. Although honey bees are often viewed as ineffective pollinators of blueberries, they successfully move pollen between flowers and often increase blueberry fruit production (Aras et al., 1996b; Mackenzie, 1997; Sampson and Cane, 2000; Hoffman et al., 2018; Martin et al., 2021). Accordingly, we hypothesize that honey bees considerably increase fruit yield and decrease fruit developmental period, compared to fruit produced in the absence of pollinators. Additionally, we establish if the pollination environment could be optimized so that fruit yield is improved relative to the present conditions where honey bees are the sole pollinators. Despite honey bees moving pollen between blueberry flowers (Hoffman et al., 2018), we predict considerable room for improvement, with hand pollination producing substantially larger fruit yields than honey bee pollination.

## 2. Methods

An experimental plot on Backsberg blueberry farm was used as the study site (Western Cape, South Africa, 33°48′30.7″S 18°54′09.8″E). Farmlands surround the experimental plot and there is also a nearby (2.98km) nature reserve consisting of Swartland shale renosterveld and fynbos. Honey bee hives were stocked at 15 hives per hectare, which falls within the typical range of commercial blueberry farm hive densities (4 – 39.5 hives per hectare; Isaacs and Kirk, 2010; Gibbs et al., 2016; Hoffman et al., 2018). Hive densities on blueberries are often much greater than on crops such as apples and pears, where two and four honey bee hives per hectare, respectively, provide sufficient pollination (Allsopp et al., 2008). This experiment comprised three treatments: pollinator exclusion, open honey bee pollination, and hand pollination. We determined the effects of honey bee pollination on blueberry production and also whether there was scope to improve pollination conditions by comparing fruit yield (fruit mass, fruit set, and developmental period) among these three treatments.

Varieties used were Emerald (*Vaccinium corymbosum* hybrid), Eureka (*Vaccinium corymbosum* hybrid), Snowchaser (Vaccinium sp. hybrid), Suziblue (*Vaccinium corymbosum* hybrid), and Twilight (*Vaccinium corymbosum* hybrid). Pollination combinations were repeated three times per plant and across nine plants, resulting in 27 pollination combination replicates per variety. Moreover, we minimized the effects of extrinsic factors, which potentially influence fruit development, by conducting our experiments across varieties in an experimental plot of only 50 m X 115 m. To determine pollination deficits and benefits of bees to the five varieties, we followed the methods of Martin et al. (2021). Below we provide a brief description (also see Fig. 1).

**Fig 1:**
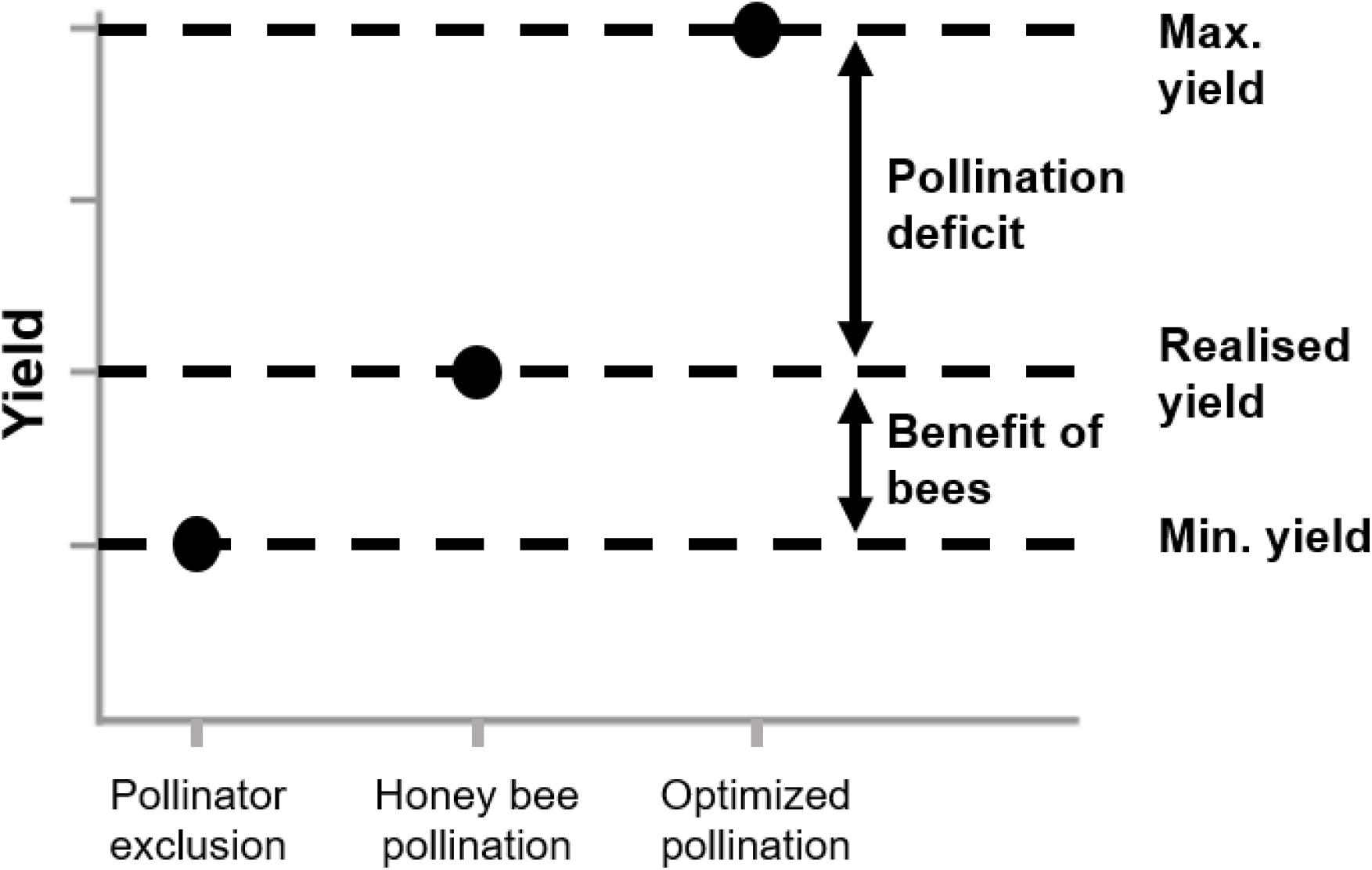
Hypothetical figure showing how three treatments can be used to determine the benefits of honey bee pollination as well as how realised yields may potentially be improved (originally produced by Martin et al. 2021).

### 2.1 Benefit of bees

We determined the benefit of bees by comparing blueberry fruit production under honey bee pollination to blueberry fruit production in the absence of honey bees (pollinator exclusion). Pollinators were denied access to flowers by covering 27 individual virgin flowers with fine mesh bags. Once flowers had wilted, these bags were removed to allow for normal fruit maturation. We estimated the fruit yield through honey bee pollination by exposing (unbagging) 27 flowers to bee pollinators. For the duration of the time spent at the experimental plot (approximately 280 hours), honey bees were the only pollinator seen visiting blueberry flowers, which is typical for South African blueberry farms.

### 2.2 Pollination deficit

We calculated the pollination deficit for each variety by comparing the honey bee treatment to the optimized hand pollination treatment. Hand pollination is thought to result in the best fruit quality as it enhances both pollen deposition and quality of pollen. Data from the optimized hand-pollination treatment were gathered at the same time as other treatments but was previously published in Martin et al. (2019). The aims of Martin et al. (2019) were exclusively to determine which varieties were most compatible, whereas the aims of this paper were to calculate the benefit of bees and pollination deficit which require data sets in addition to the ones collected by Martin et al. (2019). For the current manuscript, we used data from the best pollen donator (i.e., the cross-combination resulting in the highest yield) for each variety from Martin et al. (2019), to represent the optimized hand-pollination treatment. We followed hand pollination protocols as set out by Martin et al. (2021).

### 2.3 Measurements of fruit quality

Following pollination treatments, fruit development was examined weekly to determine whether fruit were ready to be picked. Fruits were considered ready for picking when the entire berry had transformed to a uniform dark blue. Fruit were then harvested and the weight, diameter, and developmental period (weeks from pollination to harvesting) were determined on the harvest day. We also calculated the percentage fruit set per treatment. Fruit mass and percentage fruit set were combined to calculate the adjusted fruit mass for each treatment as follows:

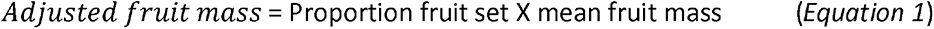

### 2.4 Statistical analyses

Linear mixed-effects models were used with fruit mass, developmental period, and adjusted fruit mass as functions of pollination treatment (fixed effect). To account for variation between individual plants, plant ID was used as a random effect. This increased the sensitivity of the model in identifying differences in fruit yield measures as a consequence of treatment alone. Linear contrasts (Tukey) were used to detect treatment differences. Nakagawa *R^2^* values (Nakagawa and Schielzeth, 2013) were used to assess model fit, providing conditional variance (*R^2^c*) and marginal variance (*R^2^m*) estimates for linear mixed-effects models, which can be equated to traditional *R^2^* values. The conditional *R^2^* values (*R^2^c*) explain the variance of the entire model (fixed effects and random effects), whereas marginal *R^2^* values (*R^2^m*) explain the variance of the fixed effects alone.

All statistical analyses were conducted in R (version 3.3.2) (R Core Team, 2017) using the packages nlme (Pinheiro et al., 2017), multcomp (Hothorn et al., 2008), ggplot2 (Wickham, 2009), sjPlot (Lüdecke, 2017), car (Fox and Weisberg, 2011), lme4 (Bates et al., 2015), grid (R Core Team, 2017), gridExtra (Auguie, 2017), lattice (Deepayan, 2008), MuMIn (Barton, 2017), plyr (Wickham, 2011), and plotrix (Lemon, 2006).

## 3. Results

### 3.1 Emerald

#### 3.1.1 Benefit of bees

We found strong statistical support indicating that honey bees increased blueberry production in Emerald compared to fruits produced in the absence of pollinators for three-quarters of the metrics measured. We provide further details on the four metrics below.

The conditional variance for the developmental period model was *R^2^c*=0.471 and the marginal variance was *R^2^m*=0.062. This shows that variation was mostly explained by the random effect (plant ID). The presence of honey bee pollinators appeared to decrease (statistically borderline) the developmental period of blueberry fruits by about 2 weeks (*z*=- 2.085, *p*=0.089, Fig 2) compared to when they were absent.

**Fig 2:**
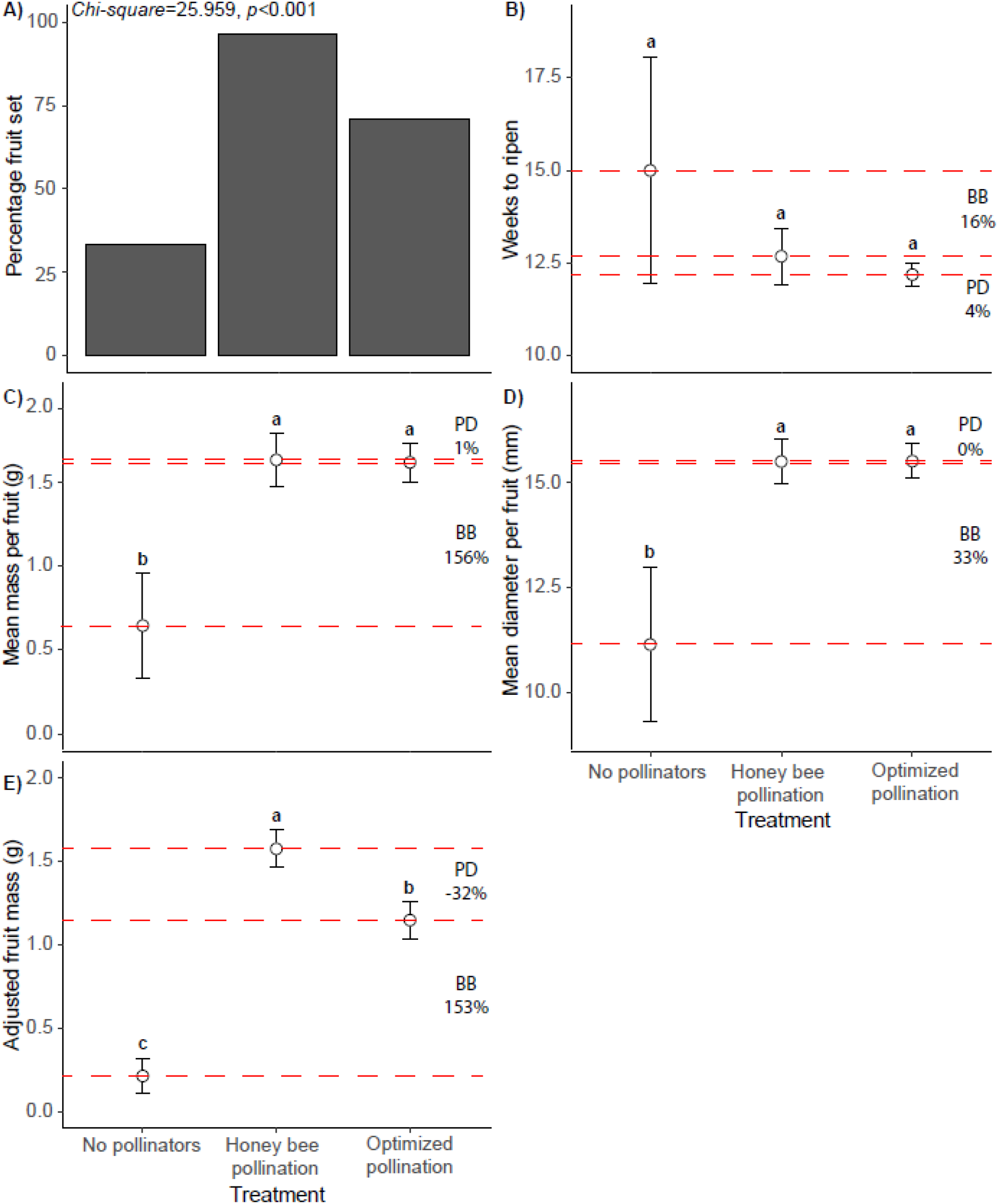
Pollination deficit and the benefit of bees for Emerald fruit. A) Percentage fruit set. B) Developmental period. C) Mean fruit mass. D) Mean fruit diameter. E) Adjusted fruit mass. Significance (p<0.05) of linear contrasts (Tukey HSD) are indicated by letters. Error bars indicate standard error. BB = benefit of bees, the difference (in percentage) between fruit produced in the absence of pollinators compared (n=27) to fruits produced by honey bees (n=27). PD = pollination deficit, the difference (in percentage) between fruits produced after honey bee exposure compared to fruits produced after hand pollination (n=27).

The conditional variance for the Emerald fruit mass model was *R^2^c*=0.238 while the marginal variance was *R^2^m*=0.151. The conditional variance for the fruit diameter model was *R^2^c*=0.303 and the marginal variance was R m=0.210. This suggests that for both fruit mass and diameter, variation is primarily explained by treatment. Fruit mass was significantly increased by the presence of honey bees (mean ± SE fruit mass = 1.64g ± 0.12g, *z*=2.799, *p*=0.013), resulting in a 156% mass increase compared to fruit produced in the absence of pollinators (mean ± SE fruit mass = 0.64g ± 0.32g, Fig 2C). Likewise, honey bees increased the fruit diameter by 4mm (33%) per fruit (mean ± SE = 15.51mm ± 0.41mm) compared to treatments produced in the absence of pollinators (mean ± SE = 11.12mm ± 1.86mm, Fig 2D, *z*=3.412, *p*=0.002). There were significant differences in fruit set following the three treatments (*Chi-square*=25.959, *df*=2, *p*<0.001, Fig 2A). Honey bees increased fruit set by 63% relative to bagged plants. Similarly, honey bee pollination outperformed the hand pollination treatment by 25%.

The conditional variance for the adjusted Emerald fruit mass model was *R^2^c*=0.478 and marginal variance *was R^2^m*=0.347. This suggests that adjusted fruit mass variation was primarily explained by treatment. Honey bees led to a significant increase of adjusted fruit mass (mean ± SE = 1.58g ± 0.11g, Fig 2E) of 153% compared to bagged treatments (mean ± SE = 0.21g ± 0.11g, Fig 2E, z=4.639, p<0.001).

#### 3.1.2 Pollination deficit

The pollination deficit metric indicated that for Emerald, honey bee pollinators performed equally or better than hand pollinations. Below we present those results.

Hand pollination resulted in a statistically non-significant (*z*=-0.073, *p*=0.997) decrease in the developmental period of blueberry fruit, from 12.68 ± 0.32 weeks (mean ± SE) when open to honey bee pollination to 12.19 ± 0.76 weeks (mean ± SE) when hand-pollinated (Fig 2B). Mass of fruit produced from hand pollination (1.62g ± 0.16g, mean ± SE), and fruit produced from honey bee pollination (1.64g ± 0.12g, mean ± SE, Fig 2C) did not differ significantly (*z*=-0.110, *p*=0.993). The diameter of hand-pollinated fruit (mean ± SE = 15.52mm ± 0.54mm) and fruit pollinated by honey bees (mean ± SE = 15.51mm ± 0.41mm), also did not differ significantly (*z*=-0.026, *p*=0.999). Honey bees outperformed hand pollination with regards to fruit set by 25%.

The adjusted fruit mass decreased by 32% (*z*=-2.471, *p*=0.034) when hand-pollinated (mean ± SE = 1.15g ± 0.11g, Fig 2E) in comparison to fruit produced from honey bee pollination (mean ± SE = 1.58g ± 0.11g, Fig 2E).

### 3.2 Eureka

#### 3.2.1 Benefit of bees

For all four metrics measured we found no statistical evidence suggesting that honey bees improved Eureka fruit quality compared to fruits produced in the absence of pollinators. We provide further details on each metric below.

For the developmental period model, the conditional variance was *R^2^c*=0.405 and the marginal variance was R m=0.370. The variation is therefore explained mostly by the fixed effect (treatment). Honey bees did not change the developmental period (*z*=-0.042, *p*=0.999, Fig 3B).

**Fig 3:**
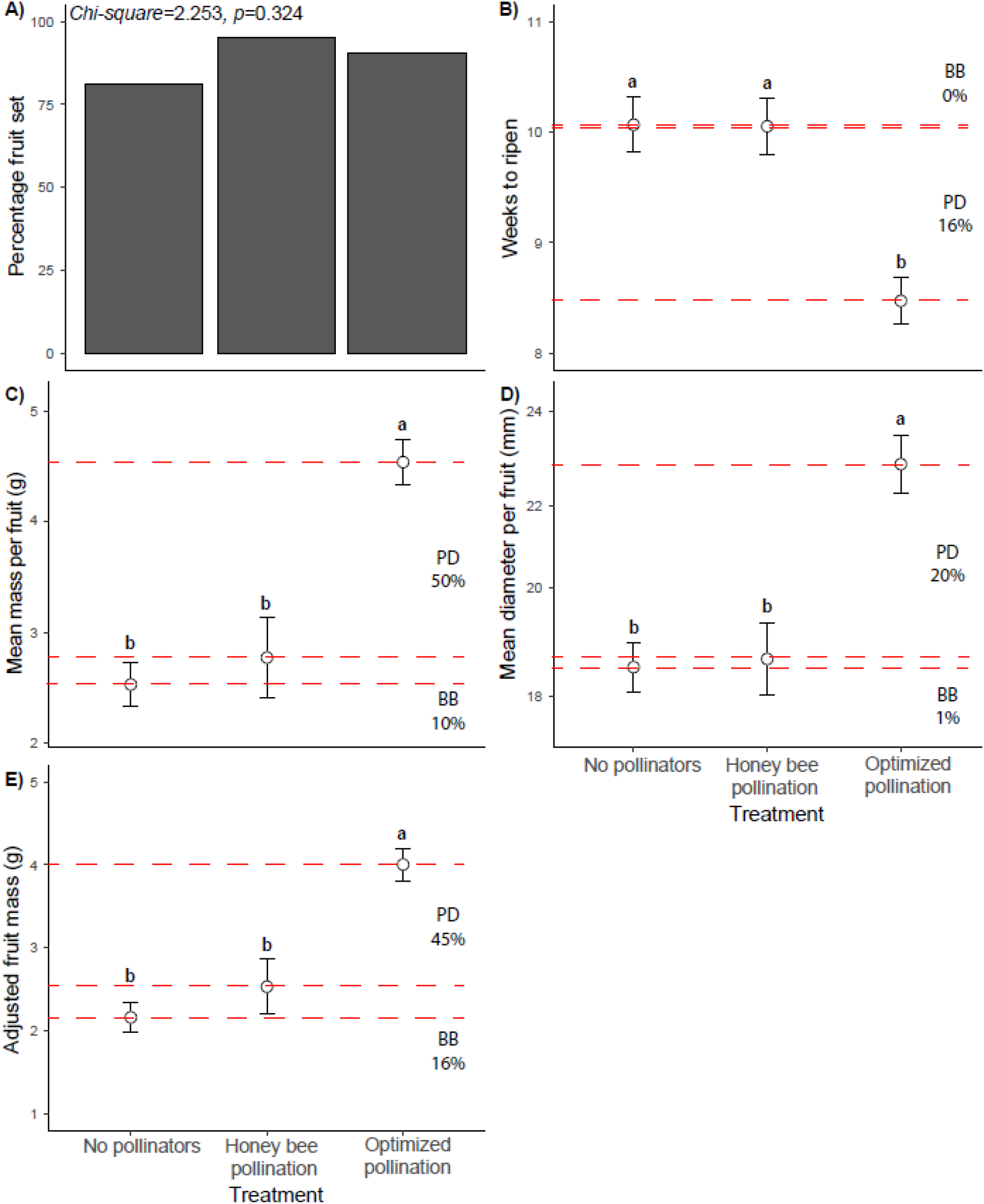
Pollination deficit and the benefit of bees for Eureka fruit. A) Percentage fruit set. B) Developmental period. C) Mean fruit mass. D) Mean fruit diameter. E) Adjusted fruit mass. Significance (p<0.05) of linear contrasts (Tukey HSD) are indicated by letters. Error bars indicate standard error. BB = benefit of bees, the difference (in percentage) between fruit produced in the absence of pollinators compared (n=27) to fruits produced by honey bees (n=27). PD = pollination deficit, the difference (in percentage) between fruits produced after honey bee exposure compared to fruits produced after hand pollination (n=27).

The fruit mass model resulted in a conditional variance of *R^2^c*=0.693 and a marginal variance of R m=0.398. The conditional variance was *R^2^c*=0.569 for the fruit diameter model, while the marginal variance was *R^2^m*=0.371. This indicated that variance is primarily explained by treatment for fruit mass and diameter. Honey bee presence did not increase fruit mass (mean ± SE fruit mass = 2.66g ± 0.21g) significantly (*z*=0.815, *p*=0.684), compared to when pollinators were absent (mean ± SE fruit mass = 2.41g ± 0.20g, Fig 3C). Likewise, honey bees did not significantly increase (*z*=0.232, *p*=0.970) fruit diameter (mean ± SE = 18.50mm ± 0.61mm) compared to when they were absent (mean ± SE = 18.33mm ± 0.52mm, Fig 3D,). Honey bee pollination appeared to increase fruit set by 14%, however, the differences between the three treatments were not significantly different (*Chi-square*=2.253, *df*=2, *p*=0.324, Fig 3A).

The conditional variance for Eureka adjusted fruit mass was *R^2^c*=0.678 and the marginal variance was R m=0.383. This indicated that for the adjusted fruit mass model, most of the variance was explained by pollination treatment. The adjusted fruit mass of honey bee pollination (mean ± SE = 2.53g ± 0.20g, Fig 3E), did not differ significantly (*z*=1.334, *p*=0.363) from the adjusted fruit mass of fruit produced in the absence of honey bees (mean ± SE = 2.16g ± 0.18g, Fig 3E).

#### 3.2.2 Pollination deficit

For all pollination deficit metrics measured for Eureka, hand pollinations outperformed honey bee pollination. We provide further details on each metric below.

The developmental period of fruit produced by hand pollination was significantly decreased by approximately one week (*z*=-4.521, *p*<0.001), from 10.05 ± 0.26 weeks (mean ± SE) when honey bee pollinated to 8.47 ± 0.26 weeks (mean ± SE) when hand-pollinated (Fig 3B). Hand pollination significantly increased (*z*=3.239, *p*=0.003) the mass of fruit by 50% (4.42g ± 0.36g, mean ± SE), compared to the fruit produced by flowers exposed to honey bees (2.66g ± 0.21g, mean ± SE, Fig 3C). Furthermore, hand pollination also significantly increased (z=3.657, p<0.001) fruit diameter (mean ± SE = 22.61mm ± 0.76mm) by 20% in comparison to honey bee pollinated flowers (mean ± SE = 18.50mm ± 0.61mm, Fig 3D). Hand pollination did not increase fruit set compared to honey bee pollination and in fact, led to a 4% decrease in fruit set.

The adjusted fruit mass increased significantly (*z*=3.008, *p*<0.007) by 45% after being hand-pollinated (mean ± SE = 4.00g ± 0.33g, Fig 2E), compared to the adjusted fruit mass after honey bee exposure (mean ± SE = 2.53g ± 0.20g, Fig 3E).

### 3.3 Snowchaser

#### 3.3.1 Benefit of bees

For all four metrics, access by honey bees substantially improved Snowchaser fruit production compared to when they were excluded. We provide further details on each metric below.

The conditional variance of the Snowchaser developmental period model was *R^2^c*=0.482, while the marginal variance was R m=0.460, suggesting that the variance in the developmental period was explained primarily by treatment (fixed effect). Honey bee pollinators significantly shortened the developmental period of blueberry fruits (mean ± SE fruit weeks = 11.48 weeks ± 0.20 weeks, *z*=-3.533, *p*=0.001, Fig 4B) by approximately two weeks compared to fruits produced in the absence of honey bees (mean ± SE fruit weeks = 13.14 weeks ± 0.80 weeks, Fig 4B).

**Fig 4:**
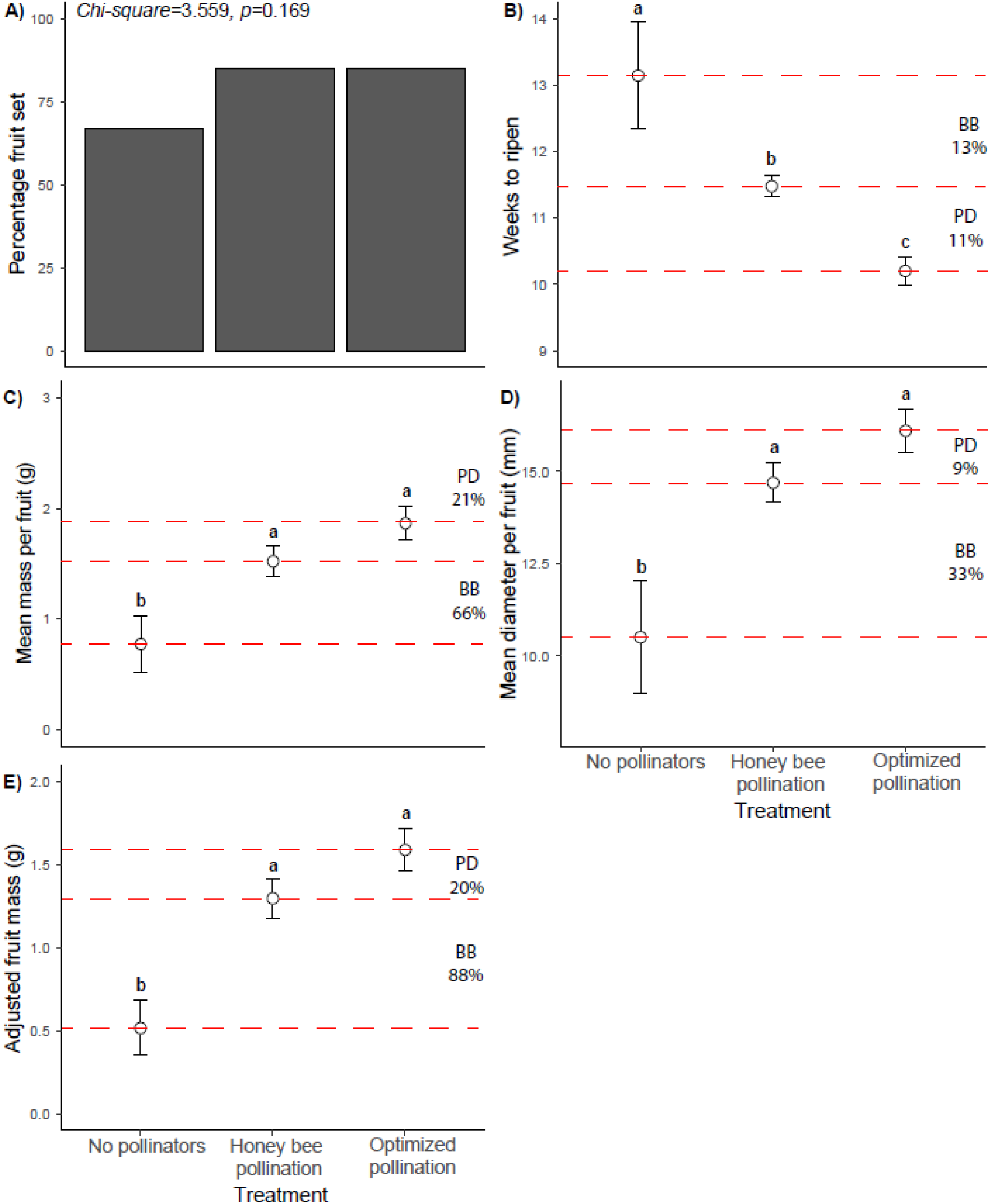
Pollination deficit and the benefit of bees for Snowchaser fruit. A) Percentage fruit set. B) Developmental period. C) Mean fruit mass. D) Mean fruit diameter. E) Adjusted fruit mass. Significance (p<0.05) of linear contrasts (Tukey HSD) are indicated by letters. Error bars indicate standard error. BB = benefit of bees, the difference (in percentage) between fruit produced in the absence of pollinators compared (n=27) to fruits produced by honey bees (n=27). PD = pollination deficit, the difference (in percentage) between fruits produced after honey bee exposure compared to fruits produced after hand pollination (n=27).

The conditional variance of the fruit mass model was R c=0.520 and the marginal variance was *R^2^m*=0.203. Similarly, the conditional variance of the fruit diameter model was *R^2^c*=0.469, while the marginal variance was R m=0.284. This suggests that variance in Snowchaser fruit mass and diameter was primarily explained by treatment. Here, honey bee presence significantly increased fruit mass (mean ± SE fruit mass = 1.52g ± 0.15g, *z*=2.362, *p*=0.047), by 66% compared to treatments where honey bees were excluded (mean ± SE fruit mass = 0.77g ± 0.25g, Fig 4C). Likewise, honey bee exposure resulted in a 4mm (33%) mean increase in fruit diameter per fruit (mean ± SE = 14.70mm ± 0.59mm) in comparison to fruit produced in the absence of honey bees (mean ± SE = 10.51mm ± 0.81mm, Fig 4D, *z*=3.229, *p*=0.003). Honey bee pollination appeared to increase fruit set by 18%, however, the differences between the three treatments were not significantly different (*Chi-square*=3.559, *df*=2, *p*=0.169, Fig 4A).

The adjusted fruit mass model resulted in a conditional variance of *R^2^c*=0.576 and a marginal variance of R m=0.273, suggesting that most of the variance in adjusted fruit mass is explained by treatment. Honey bee pollination significantly (*z*=3.062, *p*=0.006) increased adjusted fruit mass (mean ± SE = 1.30g ± 0.13g, Fig 4E), by 86% compared to the adjusted fruit mass of treatments where pollinators were excluded (mean ± SE = 0.52g ± 0.17g, Fig 4E).

#### 3.3.2 Pollination deficit

Statistics indicated that honey bee pollinators and hand pollinations performed similarly for Snowchaser in three of the four metrics measured. We provide further details below.

Hand pollination resulted in a one week or 11% decrease in the developmental period (*z*=-3.788, *p*<0.001), from 11.48 ± 0.20 weeks (mean ± SE) after honey bee exposure to 10.20 ± 0.16 weeks (mean ± SE) after hand pollination (Fig 4B). Hand-pollinated fruits had similar fruit mass (mean ± SE fruit mass = 1.87g ± 0.14g, *z*=1.469, *p*=0.303) to fruit produced after exposure to honey bees (1.52g ± 0.15g, mean ± SE, Fig 4C). Hand pollination slightly, but not significantly (z=1.425, p=0.325) increased fruit diameter (mean ± SE = 16.10mm ± 0.54mm) by 9% in comparison to flowers exposed to honey bees (mean ± SE = 14.70mm ± 0.59mm, Fig 4D). Hand pollination did not lead to an increase in fruit set compared to honey bee pollination.

The adjusted fruit mass appeared to increase by 20% for hand-pollinated flowers (mean ± SE = 1.59g ± 0.12g, Fig 2E) compared to the adjusted fruit mass after honey bee exposure (mean ± SE = 1.30g ± 0.13g, Fig 4E), however, this difference was not statistically significant (*z*=1.486, *p*=0.295).

### 3.4 Suziblue

#### 3.4.1 Benefit of bees

We found evidence suggesting that for Suziblue, honey bees do not improve blueberry fruit quality compared to when pollinators were excluded. We provide further details on each metric below.

Conditional variance for the developmental period model was *R^2^c*=0.235 and the marginal variance was *R^2^m*=0.235, suggesting that the variation was mostly explained by the fixed effect (treatment). Exposure to honey bees did not significantly affect the developmental period of Suziblue fruits (*z*=-0.957, *p*=0.604, Fig 5B).

**Fig 5:**
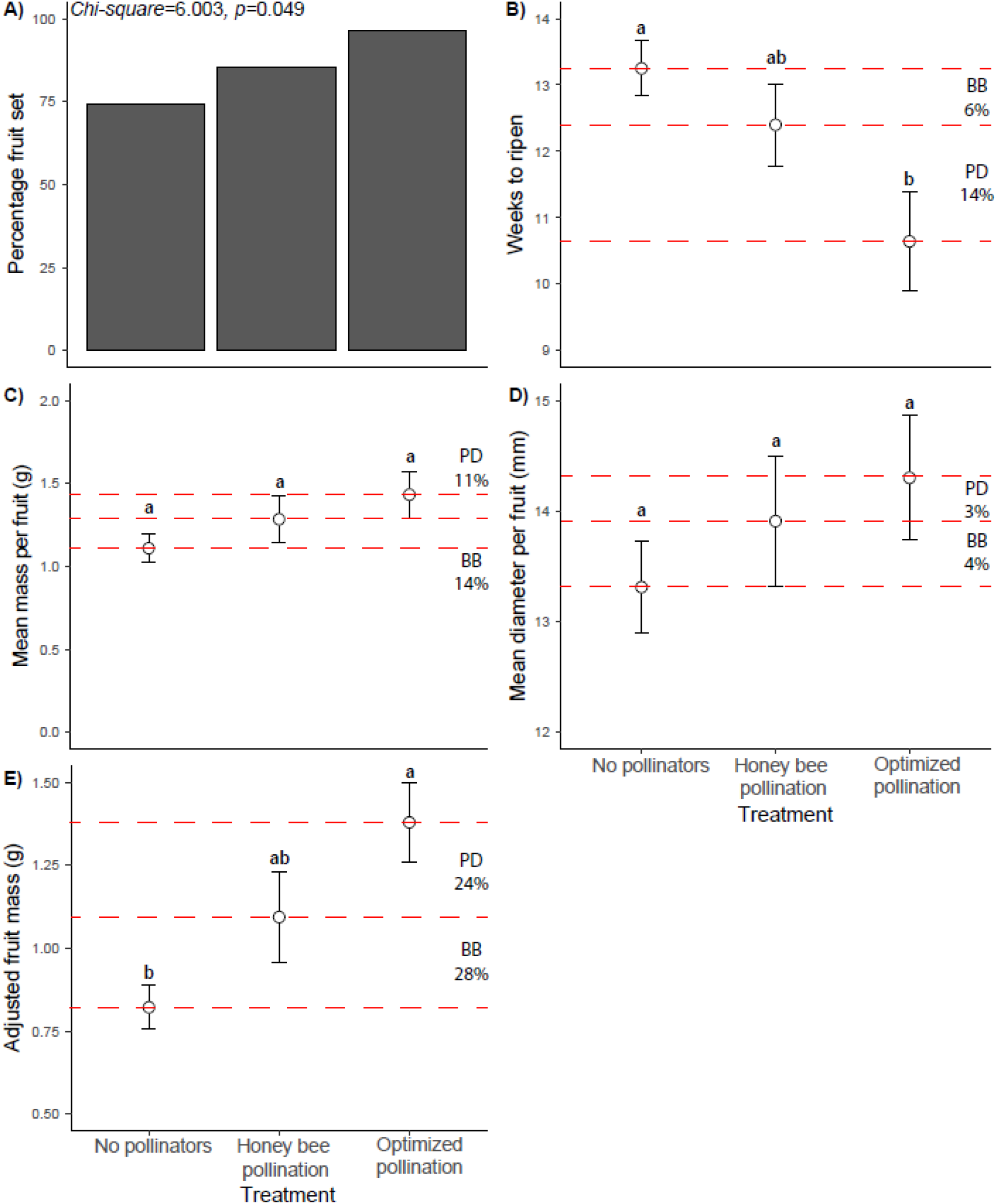
Pollination deficit and the benefit of bees for Suziblue fruit. A) Percentage fruit set. B) Developmental period. C) Mean fruit mass. D) Mean fruit diameter. E) Adjusted fruit mass. Significance (p<0.05) of linear contrasts (Tukey HSD) are indicated by letters. Error bars indicate standard error. BB = benefit of bees, the difference (in percentage) between fruit produced in the absence of pollinators compared (n=27) to fruits produced by honey bees (n=27). PD = pollination deficit, the difference (in percentage) between fruits produced after honey bee exposure compared to fruits produced after hand pollination (n=27).

The conditional variance of the fruit mass model was *R^2^c*=0.645 and the marginal variance was *R^2^m*=0.144. For the fruit diameter model, the conditional variance was *R^2^c*=0.492 and the marginal variance was R m=0.088. Plant ID explained most of the variance for fruit mass and diameter. Honey bee pollination had a minimal effect on fruit mass (mean ± SE fruit mass = 1.28g ± 0.14g, z=1.137, p=0.485), in comparison to bagged treatments (mean ± SE fruit mass = 1.11g ± 0.09g, Fig 5C). Honey bees also had a minimal effect on fruit diameter (mean ± SE = 13.91mm ± 0.56mm) in comparison to bagged treatments (mean ± SE = 13.31mm ± 0.42mm, Fig 5D, z=1.018, p=0.562). Significant differences were found in fruit set following the three treatments (Chi-square=6.003, df=2, p=0.049, Fig 5A). Honey bees increased fruit set by 11% relative to bagged plants, however, this still left a pollination deficit of 11% (honey bee treatment compared to hand pollination).

The adjusted fruit mass model resulted in a conditional variance of *R^2^c*=0.757 and a marginal variance of *R^2^m*=0.330. The variance in the adjusted fruit mass model was explained mostly by treatment (fixed effect). Flowers open to honey bee pollination resulted in similar adjusted fruit masses (mean ± SE fruit mass = 1.09g ± 0.12g, z=1.816, p=0.160), compared to honey bee exclusion treatments (mean ± SE = 0.82g ± 0.07g, Fig 5E).

#### 3.4.2 Pollination deficit

Below, we provide details showing that for all metrics measured, hand pollinations of Suziblue flowers did not outperform honey bee pollination.

The developmental period of hand-pollinated fruit was similar (mean ± SE weeks = 10.64 ± 0.62 weeks, z=-2.047, p=0.101) to that of fruit produced from honey bee pollination (12.40 ± 0.75, mean ± SE, Fig 5B). The fruit mass of hand-pollinated fruits was also similar (mean ± SE fruit mass = 1.43g ± 0.14g, z=0.786, p=0.707), to fruit produced from honey bee pollinated flowers (1.28g ± 0.14g, mean ± SE, Fig 5C). Hand pollination did not increase (z=0.579, p=0.830) fruit diameter (mean ± SE = 14.30mm ± 0.59mm) in comparison to the diameter of fruit produced by honey bee pollination (mean ± SE = 13.91mm ± 0.71mm, Fig 5D).

Adjusted fruit mass was similar and not significantly different (z=1.48, p=0.290) for hand-pollinated flowers (mean ± SE = 1.38g ± 0.14g, Fig 5E) and for flowers exposed to honey bees (mean ± SE = 1.09g ± 0.12g, Fig 5E).

### 3.5 Twilight

#### 3.5.1 Benefit of bees

For three out of the four metrics measures, honey bee presence enhanced blueberry production compared to when pollinators were excluded. We provide further details on each metric below.

The developmental period model had a conditional variance of *R^2^c*=0.360 and a marginal variance of *R^2^m*=0.320. The fixed effect (treatment) explained most of the variation. Honey bee pollination decreased the developmental period (mean ± SE weeks = 11.29 weeks ± 0.43 weeks, Fig 6B) of blueberry fruits by approximately one week or 10% compared to fruits produced in the absence of honey bees (mean ± SE fruit weeks = 12.55 weeks ± 0.49 weeks, Fig 2C). This was borderline-significant (z=-2.170, p=0.076, Fig 6B).

**Fig 6:**
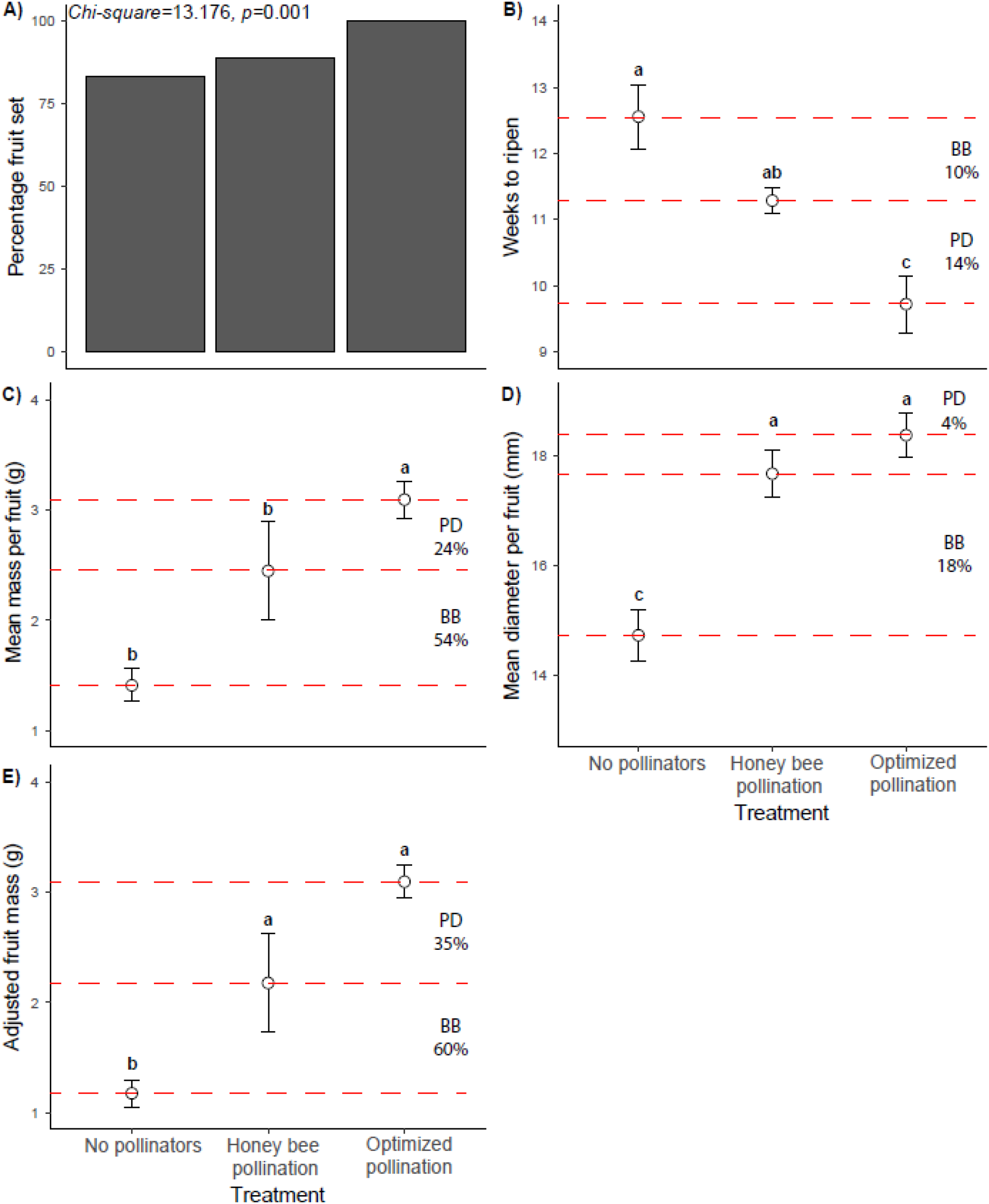
Pollination deficit and benefit of bees for Twilight fruit. A) Percentage fruit set. B) Developmental period. C) Mean fruit mass. D) Mean fruit diameter. E) Adjusted fruit mass. Significance (p<0.05) of linear contrasts (Tukey HSD) are indicated by letters. Error bars indicate standard error. BB = benefit of bees, the difference (in percentage) between fruit produced in the absence of pollinators compared (n=27) to fruits produced by honey bees (n=27). PD = pollination deficit, the difference (in percentage) between fruits produced after honey bee exposure compared to fruits produced after hand pollination (n=27).

The conditional variance for the Twilight fruit mass model was *R^2^c*=0.337 and the marginal variance was R m=0.170. For the fruit diameter model, the conditional variance was *R^2^c*=0.392, while the marginal variance was R m=0.369. This suggests that for both fruit mass and diameter, the majority of the variance was explained by treatment. Fruit mass was significantly increased after honey bee exposure (mean ± SE fruit mass = 2.45g ± 0.17g, z=2.397, p=0.043), resulting in a 54% increase compared to bagged treatments (mean ± SE fruit mass = 1.41g ± 0.17g, Fig 6C). Honey bee exposure similarly increased the mean fruit diameter by 3mm (18%) per fruit (mean ± SE = 17.66mm ± 0.39mm) in comparison to bagged treatments (mean ± SE = 14.73mm ± 0.47mm, Fig 6D, z=4.643, p<0.001). There were significant differences in fruit set following the three treatments (Chi-square=13.176, df=2, p=0.001, Fig 6A). Honey bees increased fruit set by 6% relative to bagged plants, however, this still left a pollination deficit of 11% (honey bee treatment compared to hand pollination).

The adjusted fruit mass model resulted in a conditional variance of *R^2^c*=0.383 with the marginal variance of *R^2^m*=0.216 suggesting that fruit mass variance was primarily explained by treatment. Honey bee presence significantly (z=2.368, p=0.046) increased adjusted fruit mass (mean ± SE = 2.18g ± 0.15g, Fig 6E), resulting in a 60% increase compared to pollinator exclusion treatments (mean ± SE = 1.18g ± 0.12g, Fig 6E).

#### 3.5.2 Pollination deficit

Hand pollinations resulted in improved Twilight fruit quality compared to honey bee pollinations. However, these were only statistically supported for developmental period. Below, we explain the details relating to each of the metrics measured.

The developmental period of blueberry fruit was significantly decreased by approximately one week or 14% (z=-2.824, p=0.0132), from 11.29 ± 0.43 weeks (mean ± SE) when flowers were open to honey bee pollination to 9.72 ± 0.22 weeks (mean ± SE) when flowers were pollinated by hand (Fig 6B). Hand-pollinated fruits appeared to be about 10% heavier (2.69g ± 0.13g, mean ± SE), than fruit produced from flowers exposed to honey bees (2.45g ± 0.17g, mean ± SE, Fig 6C), however, this relationship was non-significant (z=1.134, p=0.493). Hand pollination did not significantly increase (z=1.134, p=0.493) the fruit diameter (mean ± SE = 18.36mm ± 0.43mm) when compared to fruit produced by honey bee pollinated flowers (mean ± SE = 17.66mm ± 0.39mm, Fig 6D).

Hand pollination appeared to improve the adjusted fruit mass by 23% (mean ± SE = 2.69g ± 0.13g, Fig 2E) when compared to flowers exposed to honey bees (mean ± SE = 2.18g ± 0.15g, Fig 2E), although this difference was not statistically significant (z=1.694, p=0.205).

For discussion purposes, we have included a summary table which enables visual comparisons of bee benefit and pollen deficits to be made between different varieties. We have chosen to use adjusted fruit mass as a summary metric (Table 1) because it incorporates arguably the two most important yield metrics (fruit mass and fruit set). It also indirectly incorporates fruit diameter, which is highly correlated with fruit mass (Jorquera-fontena et al., 2017). Similarly, we also summarize treatment effects (figure 7) for all blueberry varieties, using adjusted fruit mass as a metric.

**Table 1:**
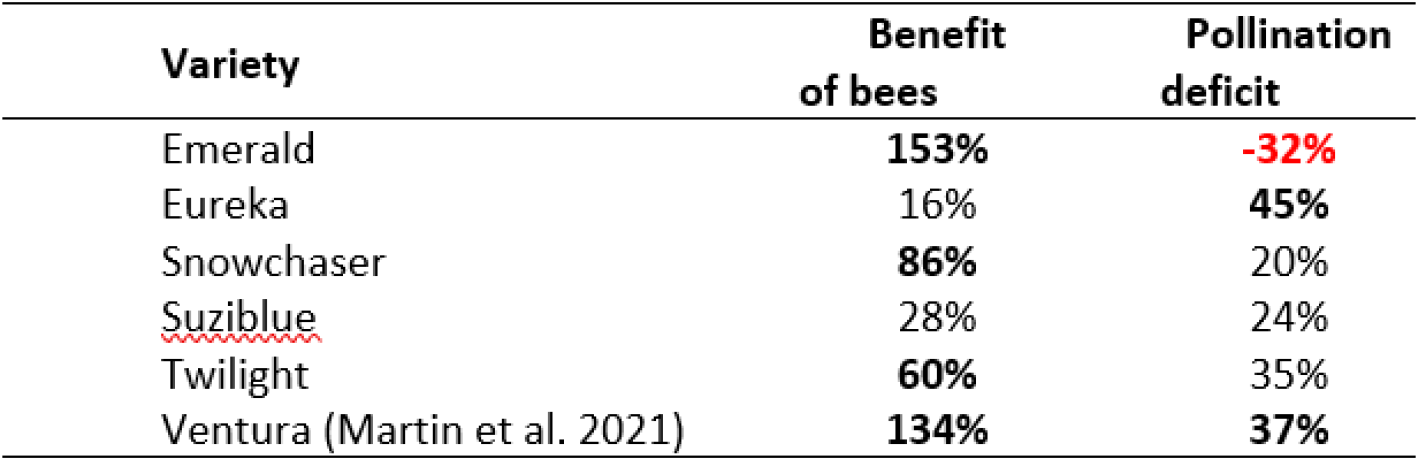
Summary table comparing the benefit of bees and the pollination deficit for adjusting fruit mass in the five blueberry varieties from this study and Ventura, which was published by Martin et al. (2021). Positive benefit of bees implies that the addition of honey bees increases fruit yields compared to when pollinators are absent. Positive pollination deficit indicates that plants pollinated by honey bees produce lower yields than plants pollinated by hand. Statistical significance (p < 0.05) is indicated by bold fonts, while colour differences (red vs black) suggest differences in trend direction.

**Fig 7:**
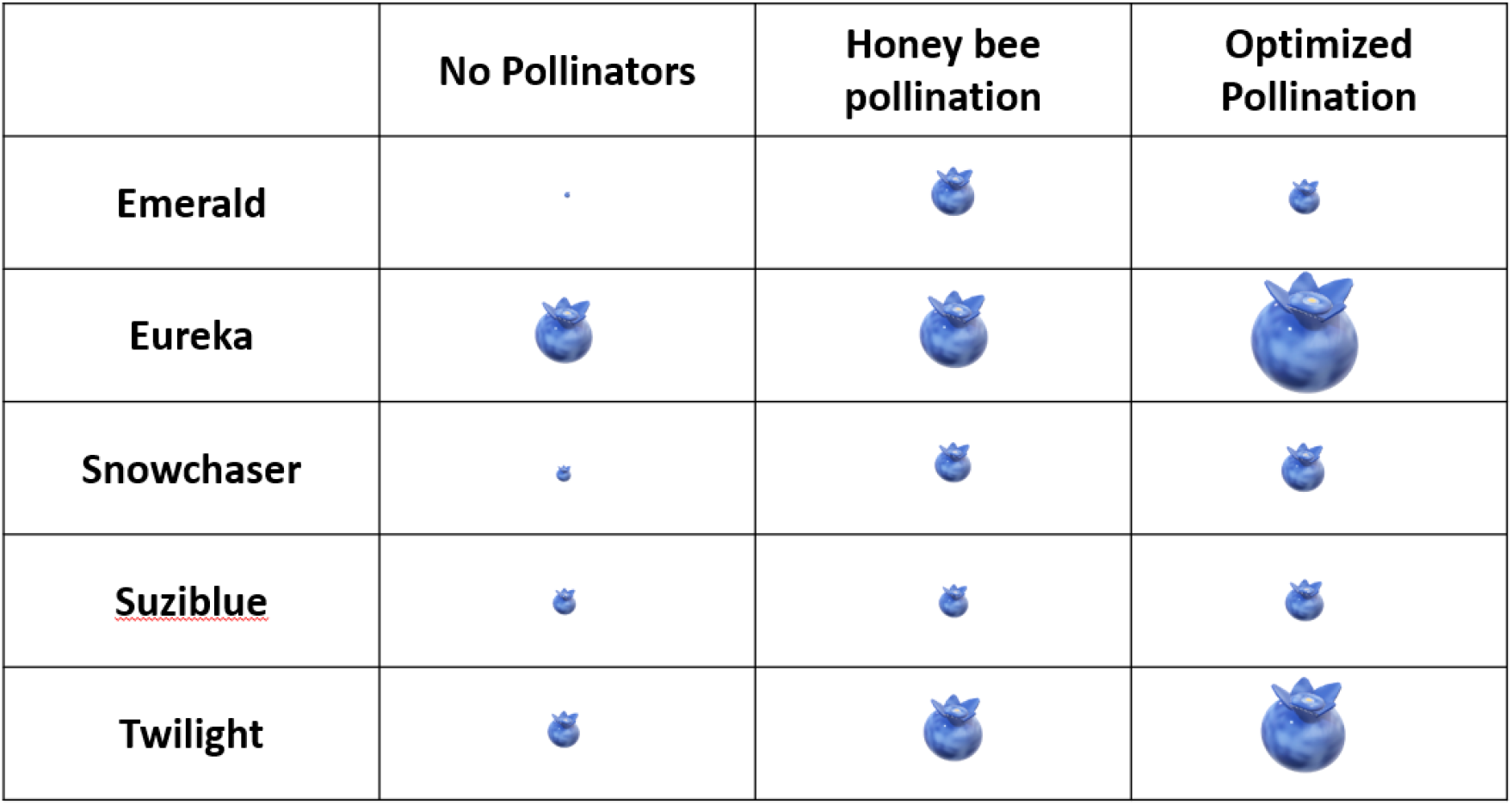
Relative size comparisons (linear scale) of treatment effects for adjusted fruit mass in the five blueberry varieties. The largest blueberry has an equivalent adjusted fruit mass of 4g while the smallest has an equivalent adjusted fruit mass of 0.21g.

## 4. Discussion

We provide a comprehensive comparison of honey bee effectiveness as pollinators of common commercial blueberry varieties in South Africa. Evidence suggests that the fruit quality of these southern highbush blueberry varieties is affected considerably by the pollination environment. Overall, we show that the absence of pollinators in highbush blueberries typically leads to reductions in fruit yield. This shows that honey bees are usually extremely beneficial and can act as reliable blueberry pollinators, even in areas without access to native blueberry pollinators. Importantly, the study also highlights differences in the effectiveness of honey bees across different blueberry varieties. Finally, it also suggests that pollination conditions could be improved for certain varieties which appear not to be producing at their full production potential under honey bee pollination.

In all cases, honey bees increase yields (usually statistically significant) compared to when pollinators were absent (Table 1). Martin et al. (2021) also demonstrated this for Ventura, another southern highbush variety used in South Africa. In this study, honey bees led to a 134% increase of adjusted fruit mass compared to fruits formed in the absence of honey bees, demonstrating that honey bees can effectively pollinate highbush blueberry varieties.

The addition of honey bees can result in very impressive yield increases for some varieties (i.e. Emerald, Snowchaser, Twilight, and Ventura), but for two varieties (Eureka and Suziblue), honey bee exposure had relatively small effects on fruit production (Fig 7). This may occur if a variety is good at autonomous self-pollination and does not need pollinators. Excellent autonomous self-pollinators should also have low pollination deficits because pollinators are not needed for good fruit production. Consequently, fruit yields without pollinators should be similar to fruit yields under optimized pollination conditions. However, none of the varieties appear to be excellent autonomous selfers and Emerald appears to be a particularly poor selfer and had minute adjusted fruit mass when pollinators were excluded. However, the pollination deficit for Emerald was remarkable because it was so small (less than zero), suggesting that honey bee pollination was more effective than hand pollinations. The presence of honey bees also had an enormous (153%) positive impact on yield, suggesting that this variety is exceptionally well suited to honey bee pollination. While the low pollination deficit of Emerald may potentially make it a good variety to use in areas with honey bee pollinators, the inherently small fruit sizes of this variety, even under optimal pollination conditions may make it less attractive than other varieties with larger pollination deficits.

Eureka, for example, had a large pollination deficit (45%), suggesting that the pollination environment was far from optimal and that improvements in pollination conditions can markedly increase yields. However, this variety has two redeeming features which make it a potentially good commercial variety, even under very poor pollination conditions. Eureka appears to be a reasonably good autonomous selfer and can set fruit in the absence of pollinators. In fact, fruit yields without pollinators are only marginally smaller (16%) than fruit yield after honey bee exposure, suggesting that honey bees do not currently add much benefit to this crop. Despite this, its adjusted fruit mass without pollinators is still larger than most other varieties when they are under optimized pollination conditions (except Twilight) (Fig 7).

Snowchaser appears to be a poor autonomous self-pollinator and had a very small adjusted fruit mass in the absence of honey bees. However, honey bee presence had a significantly positive impact on Snowchaser yields and this, in combination with small (20%) pollination deficits, suggests that this variety is also quite well suited to honey bee pollination. Unfortunately, this advantage may be outweighed by its small fruit size (even under optimized pollination conditions) which is likely to reduce its attractiveness as a commercial crop.

Suziblue appears to be a good autonomous self-pollinator because it is able to produce yields without pollinators, which are similar to the yields after honey bee exposure. This suggests that autonomous self-pollination does a reasonable job of ensuring fruit production and that yields are not strongly increased by the presence of honey bees. Furthermore, it had one of the smaller (non-significant) pollination deficits measured (24%). Although it may appear as though Suziblue is potentially a good variety to plant in conditions with unpredictable pollinator variability, its generally small fruit size, even under optimal pollination conditions leads to low adjusted fruit mass.

Twilight does not appear to produce its best fruit in the absence of pollinators, although the fruits are nevertheless still large compared to several of the other varieties. The presence of honey bees made a significant increase in Twilight yields although there still appeared to be a large (but not significant) pollination deficit. While the large pollination deficit suggests that Twilight is not well suited to the honey bee pollination environment, its inherently large fruit size enables it to produce large fruit under honey bee pollination, with plenty of potential to improve pollination. The large fruit size compared to other varieties under honey bee pollination may make this variety a good commercial candidate despite the relatively large pollination deficit.

Importantly, our interpretations of yield data in this manuscript do not consider potential differences in the number of buds produced by different varieties of blueberry plants as we were not able to access these data. Also, we did not attempt to determine the temporal value of yield produced by each variety. For example, early or late season varieties may demand higher export prices than mid-season varieties when the export market is saturated. These market-level considerations are beyond the scope of the manuscript.

It is not always clear exactly why some varieties differ so markedly in the size of their pollination deficits. However, in some cases, varieties (e.g. Suziblue) that are good at autonomous selfing can reduce the pollination deficit. In others, the pollination deficit can be reduced if varieties are well suited to honey bee pollination (e.g. Emerald). While we did not study the mechanics of “suitability to honey bee pollination,” this may depend on how well blueberry and honey bee morphology are matched. Blueberry flowers can differ significantly in terms of their morphological characters and certain floral traits may make it difficult for honey bees to access rewards. This may reduce pollinator visitation rates to those varieties or possibly make it hard for pollen grains to be extracted. Studies on wild pollinators have often found that traits such as floral tube length and diameter can have a profound influence on the effectiveness of pollen transfer and also on whether pollinators are able to access nectar or not (Campbell et al., 1996; Aigner, 2004; Fenster et al., 2006; Martins and Johnson, 2013; Lázaro et al., 2015; Minnaar et al., 2019). It is possible that varieties with long tubes can exclude pollinators from accessing nectar, especially if the tubes are also narrow and do not allow the bees to enter very deeply into the flowers. Varieties with narrow entrances may also exclude pollinators from accessing both nectar and pollen, even if the tube lengths are short. Evidence that some varieties have inaccessible or poor rewards for pollinators can be seen in studies that show strong pollinator preferences for one blueberry variety over another (Dorr and Matin, 1966; Courcelles et al., 2013).

Honey bees are often thought to be poor pollinators of blueberries because of their reluctance to forage in cold weather conditions (Tuell and Isaacs, 2010), slower foraging times (Javorek et al., 2002), and decreased pollen deposition per visit (Javorek et al., 2002) when compared to pollinators such as bumble bees. Despite this, we show that honey bees are likely to be reliable pollinators of several blueberry varieties in regions that do not have access to bumble bees. This is especially true for Emerald, where honey bee pollination is so effective that it can potentially negate the need for other pollinators. While the small fruits of this variety may make it less attractive for commercial use, its suitability to honey bee pollination suggests that its floral characteristics may hold the key to improving fruit quality of varieties still to be developed in the future. Inherent differences in fruit size among blueberry varieties may be a very important factor guiding the choice of variety. However, considerable variation in autonomous selfing ability, benefit of bees, and pollination deficits across varieties suggest that it may be possible to identify the plant traits responsible for these differences and actively select for those traits in the development of new varieties.

Considerable research is needed to identify which blueberry varieties do best under honey bee pollination under commercial conditions, what makes varieties successful or unsuccessful, which varieties can be effectively pollinated by honey bees, and which varieties need other effective means to increase fruit quality. Future research should therefore aim to determine how to decrease the pollination deficit of blueberry varieties either by increasing the effectiveness of autonomous selfing or by increasing the effectiveness of honey bees.

## Notes

### Competing Interest Statement

The authors have declared no competing interest.

